# Circulating mast cell progenitors migrate to atopic dermatitis-like skin then adapt locally

**DOI:** 10.1101/2022.03.04.483030

**Authors:** Yuki Honda Keith, Tetsuya Honda, Sachiko Ono, Bernett Lee, Sho Hanakawa, Yoshihiro Ishida, Satoshi Nakamizo, Kenji Kabashima

## Abstract

**Background:** Mast cells (MCs) are tissue-resident immune cells that are classified into two subsets in mice: connective tissue-type MCs (CTMCs) and mucosal type MCs (MMCs). Although both CTMCs and MMCs can be induced from bone marrow (BM)-derived hematopoietic stem cells (HSCs) in vitro, recent research on MC ontogeny has revealed that MMCs are maintained with a supply of BM-derived HSCs, while CTMCs are maintained locally by self-proliferation in steady state *in vivo*. However, how CTMCs, such as skin MCs, are maintained in an inflammatory state such as atopic dermatitis (AD) remains to be fully elucidated. Methods: MC903-induced AD model was used to identify BM-derived MCs in the skin. The infiltration and proliferation of MCs were evaluated by flow cytometry using CD45.1 BM-chimera mice and parabiosis. BM-derived MCs in AD-like skin were compared to resident MCs (rMCs) in gene expressions by RNA sequence analysis. The fate of BM-derived MCs in AD-like skin was investigated for expressions of CTMC markers and responses to compound 48/80. Results: In AD-like skin, significant increase of both rMCs and BM-derived MCs was observed. BM-derived MCs were derived from circulating MC progenitors (MCps) and were distinguished from rMCs by integrinβ7 expression, which was gradually downregulated in the skin. RNA sequence analysis showed that integrinβ7^+^ MCs in the skin shared characteristics of both MMC and CTMC. Integrinβ7^+^ MCs proliferated in situ and acquired the CTMC phenotypes in AD-like skin. Conclusions: Skin MCs are maintained in AD-like skin by both local proliferation of rMCs and infiltration/proliferation of BM-derived MCs, which differentiate toward CTMC in the skin.

## Introduction

Mast cells (MCs) are tissue-resident immune cells with various functions distributing throughout the tissues. Murine MCs are classified into two subsets: connective tissue-type MCs (CTMCs) and mucosal-type MCs (MMCs).(Kitamura, 1989) Both CTMCs and MMCs express high-affinity IgE receptors (FcεRI) and mediate type I hypersensitivity, but there are several differences between CTMCs and MMCs.(Gurish and Austen, 2012) a) CTMCs contain tryptases, chymases and heparin in their granules, and MMCs contain only chymases. b) CTMCs reside constitutively in connective tissues including the skin, while MMCs are distributed in mucous membranes and are supplied from circulation. c) CTMCs can degranulate in response to various cationic substances via Mas-related G protein-coupled receptor b2 (Mrgprb2),(McNeil et al., 2015) while MMCs degranulate mainly by crosslinking of FceRI by binding of cognitive antigens. d) CTMCs are long-lived, while MMCs are short-lived. Similar to murine MCs, human MCs are classified into two subsets: MC_TC_s (tryptase-positive, chymase-positive) and MC_T_s (tryptase-positive, chymase-negative). MC_TC_s are dominant in connective tissues, while MC_T_s are dominant in mucosal tissues.(Craig and Schwartz, 1989)

MC progenitors (MCps) are identified in adult murine bone marrow (BM),(Chen et al., 2005) and both CTMCs and MMCs can be induced from BM-derived hematopoietic stem cells (HSCs) *in vitro*.(Takano et al., 2008), (Granule-specific et al., 1999) However, research on ontogeny has revealed that only MMCs are maintained by the supply of BM-derived MCps, while the maintenance of CTMCs is independent from the supply of BM-derived MCps in steady state.(Gentek et al., 2018), (Li et al., 2018) In mucosal tissues under inflammatory state such as allergic reactions and infections, an increased migration of MCps into the mucosal tissues is observed.(Abonia et al., 2006), (Kasugai et al., 1995) Moreover, it has recently been demonstrated that inducible MCs, possibly derived from MCps, show transcript signatures of MMCs, and are distinct from CTMCs in murine allergic lung.(Derakhshan et al., 2021) These data indicate that BM-derived MCps migrate to mucosal tissues both in steady state and under inflammation, then differentiate into MMCs.

MCs are abundant in the skin and they are CTMCs in mice and MC_TC_s in humans. Skin MCs are increased in various skin diseases including atopic dermatitis (AD).(Murota et al., 2008) AD is one of the most common inflammatory skin diseases presenting dry skin with chronic pruritic eczema. Degranulation of skin MCs can accelerate the inflammation by evoking pruritus and secreting various inflammatory mediators in AD.(Kawakami et al., 2009) Unlike MMCs, skin MCs are maintained locally without migration of BM-derived MCps in a steady state.(Gentek et al., 2018) In addition, an in vitro study showed that human skin MCs have a potential to proliferate retaining their characteristics,(Kambe et al., 2001) suggesting MCs are increased and maintained in AD skin due to their local proliferation. In contrast, there are some in vivo studies suggesting the possible migration of MCps to the inflammatory skin, in which intravenously injected BM-derived cultured MCs (BMMCs) were detected in the skin applied with certain chemotactic lipid mediators including prostaglandin E2 (PGE2) and leukotriene B4 (LTB4).(Weller et al., 2007) In addition, it was demonstrated that BM-derived MCs are increased in 12-O-tetradecanoylphorbol-13-acetate (TPA)-induced chronic dermatitis models by using BM-chimera mice.(Weitzmann et al., 2020), (Sonoda et al., 1981) However, whether BM-derived MCps contribute to the increase of skin MCs in AD, its extent, and their fate in the skin have not been elucidated.

In this study, we demonstrated that skin MCs are increased in AD-like skin by both local proliferation and infiltration of BM-derived MCps. We characterized the infiltrating MCps in a comparison with skin-resident MCs and elucidated their adaptation to the skin by differentiating to skin type CTMCs.

## Results

### Skin MCs were increased in AD-like skin both due to local proliferation and infiltration

To confirm the increase of skin MCs under allergic inflammation, AD-like skin was induced in mice ears by topical application of MC903 for 10 days (Figure 1A), and the ear skin was subjected to flow cytometric analysis and immunohistochemical analysis on day 14 for evaluation of MCs. MC903, also called as calcipotriol, is a vitamin D3 analog that can induce AD-like skin by increasing thymic stromal lymphopoietin production from keratinocytes.(Li et al., 2006) In this AD model, the ear thickness increased over time (Figure 1B). Skin MCs were gated as Live^+^ CD45^+^ Lin (Ter119, CD3, B220, Ly-6G, siglec-F)^-^ CD11c^-^ CD11b^-^ c-kit^+^ cells by flow cytometry (Figure 1C). Avidin was used to stain skin MCs by immunohistochemistry. Flow cytometric analysis and immunohistochemical analysis showed the significant increase of skin MCs in the MC903-treated skin (AD-like skin) (Figure 1D and 1E). Enzyme immuno-sorbent assay (ELISA) revealed that stem cell factor (SCF), which is considered to be responsible for MC proliferation and differentiation,(Witte, 1990) was significantly increased in the AD-like skin (Figure S1A).

**Figure 1.**
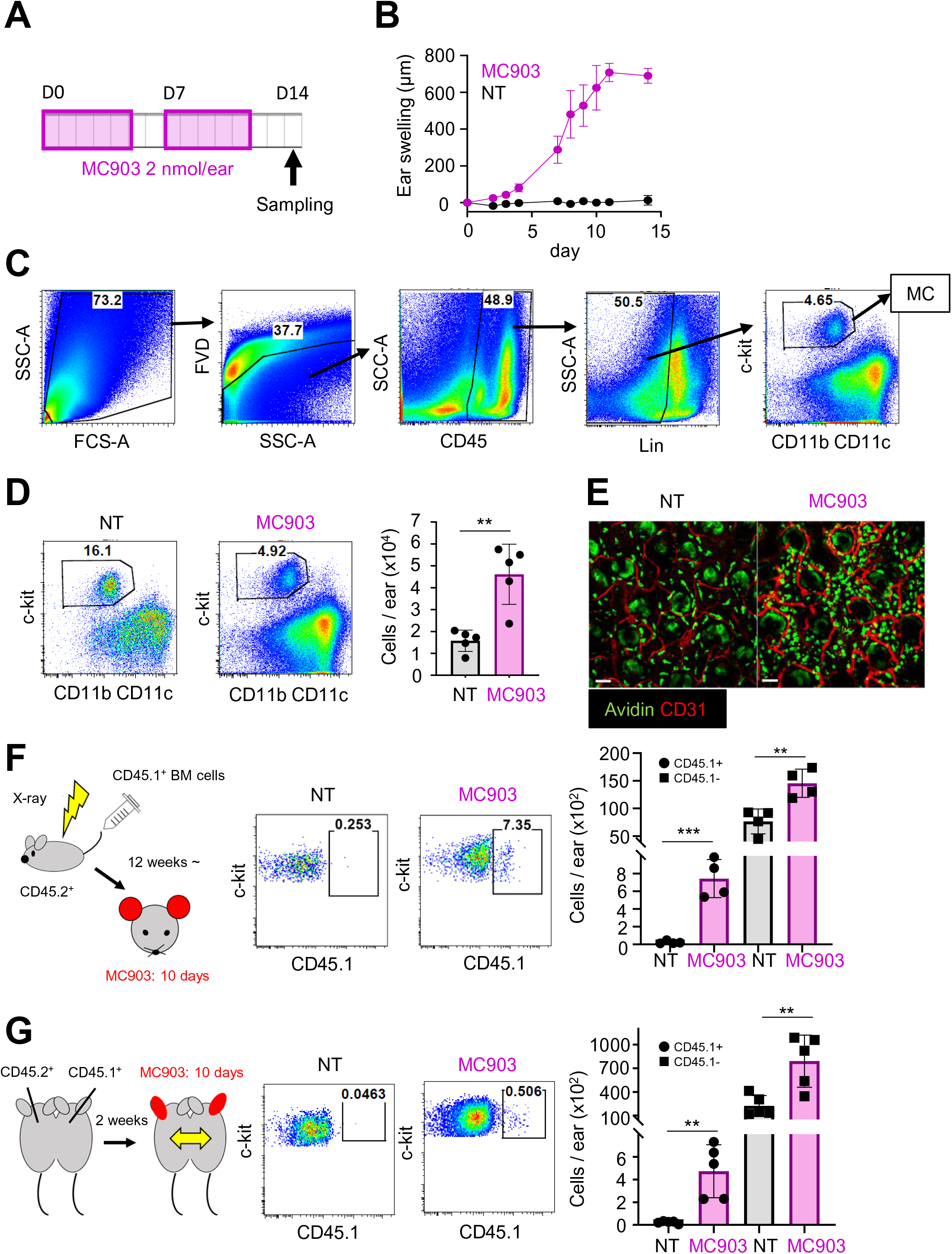
Both resident and infiltrating mast cells (MCs) were increased in MC903-induced AD-like skin. **A,** The experimental scheme of MC903-induced atopic dermatitis (AD) model. **B,** Ear swellings of non-treated (NT) and MC903-treated skin. (n = 5) **C,** Gating strategy of skin MCs. **D,** The number of skin MCs in NT and MC903-treated skin. (n = 5) **E**, Whole-mount immunostaining of avidin (green) and CD31 (red) in ear skin. Scale bar, 50 μm. **F,** Generation of CD45.1 BM-chimera mice (left panel). Representative flow cytometry plots for the expression of CD45.1 on MCs in NT or MC903-treated skin (middle panel). The number of CD45.1^+^ (circle) and CD45.1^-^ MCs (square) in NT and MC903-treated skin (right panel). (n = 4) **G,** Generation of parabiotic mice (left panel). Representative flow cytometry plots for the expression of CD45.1 on MCs in NT or MC903-treated skin (middle panel). The number of CD45.1^+^ (circle) and CD45.1^-^ MCs (square) in NT and MC903-treated skin (right panel). (n = 5) Results are expressed as the mean ± standard deviation (SD). Data are representative of at least three independent experiments. **, p < 0.01 and ***, p < 0.001.

To dissect the origin of increased skin MCs in the AD-like skin, CD45.1 BM chimera mice were generated, and were applied to the MC903-induced AD-like model 12 weeks after the BM transplantation (Figure 1F). Because skin resident MCs (rMCs) are radio-resistant,(Kitamura, 1977) rMCs and BM-derived MCs can be distinguished by the expression of CD45.1 in the chimera. Flow cytometric analysis of the ear skin revealed that there were only a few CD45.1^+^ MCs in non-treated (NT) skin, while the number of CD45.1^+^ MCs was significantly increased in the AD-like skin, suggesting that BM-derived MCs infiltrated into AD-like skin, consistent with the previous reports in TPA-induced chronic dermatitis models.(Weitzmann et al., 2020), (Sonoda et al., 1981) In addition, CD45.1^-^ MCs were also increased in the AD-like skin (Figure 1F). To further confirm the infiltration of circulating BM-derived MCs, parabiosis experiments were performed. Wild-type (WT) CD45.2^+^ mice and congenic CD45.1^+^ mice were conjoined each other, then one ear in each mouse was treated with MC903 for 10 days. The ear on the other side was left untreated (NT). Then blood and ears of WT CD45.2^+^ mice were analyzed by flow cytometry. In WT CD45.2^+^ mice, about 40% of neutrophils in blood and skin were CD45.1 positive, suggesting these mice shared their circulation successfully (Figure S1B). In NT skin in WT CD45.2^+^ mice, there were almost no CD45.1^+^ MCs, but their number was significantly increased in AD-like skin. As with the chimera experiments, CD45.1^-^ MCs were also increased in AD-like skin of WT CD45.2^+^ mice (Figure 1G). These data demonstrated that skin MCs are increased in AD-like skin both due to local proliferation of rMCs and infiltration of BM-derived MCs.

### BM-derived integrinβ7-positive MCps were identified in AD-like skin

To characterize BM-derived MCs (CD45.1^+^ MCs in MC903-treated skin) in comparison with rMCs (CD45.1^-^ MCs in NT/MC903-treated skin), the expression of common MC markers (FcεRIα, CD16/32), CTMC markers (CD200R3, heparin)(Dwyer et al., 2017), (Bergstresser et al., 1984) and MCp markers (ST-2, integrinβ7)(Dahlin et al., 2013) was analyzed by flow cytometry using CD45.1 BM chimera mice (Figure 2A). Skin MCs were pre-gated as Live^+^ CD45^+^ Lin (Ter119, CD3, B220, Ly-6G, siglec-F)^-^ CD11c^-^ CD11b^-^ c- kit^+^ cells by flow cytometry. FcεRIα and CD16/32 were expressed on all MC populations, but the expression was higher in rMCs. CD200R3 was highly expressed on rMC in both NT and AD-like skin, while the expression was partially negative on BM-derived MCs. Heparin was highly expressed in rMCs, and negative in most BM-derived MCs. ST-2 was expressed only on MCs in AD-like skin (rMCs and BM-derived MCs). Integrinβ7 was negative on rMCs, but positive on BM-derived MCs. Collectively, BM-derived MCs expressed lower levels of a CTMC signature of heparin, and higher levels of MCp signatures including integrinβ7 and ST-2. Thus, we focused on integrinβ7 as a marker of BM-derived MCs in the skin, and further examined the possibility.

**Figure 2.**
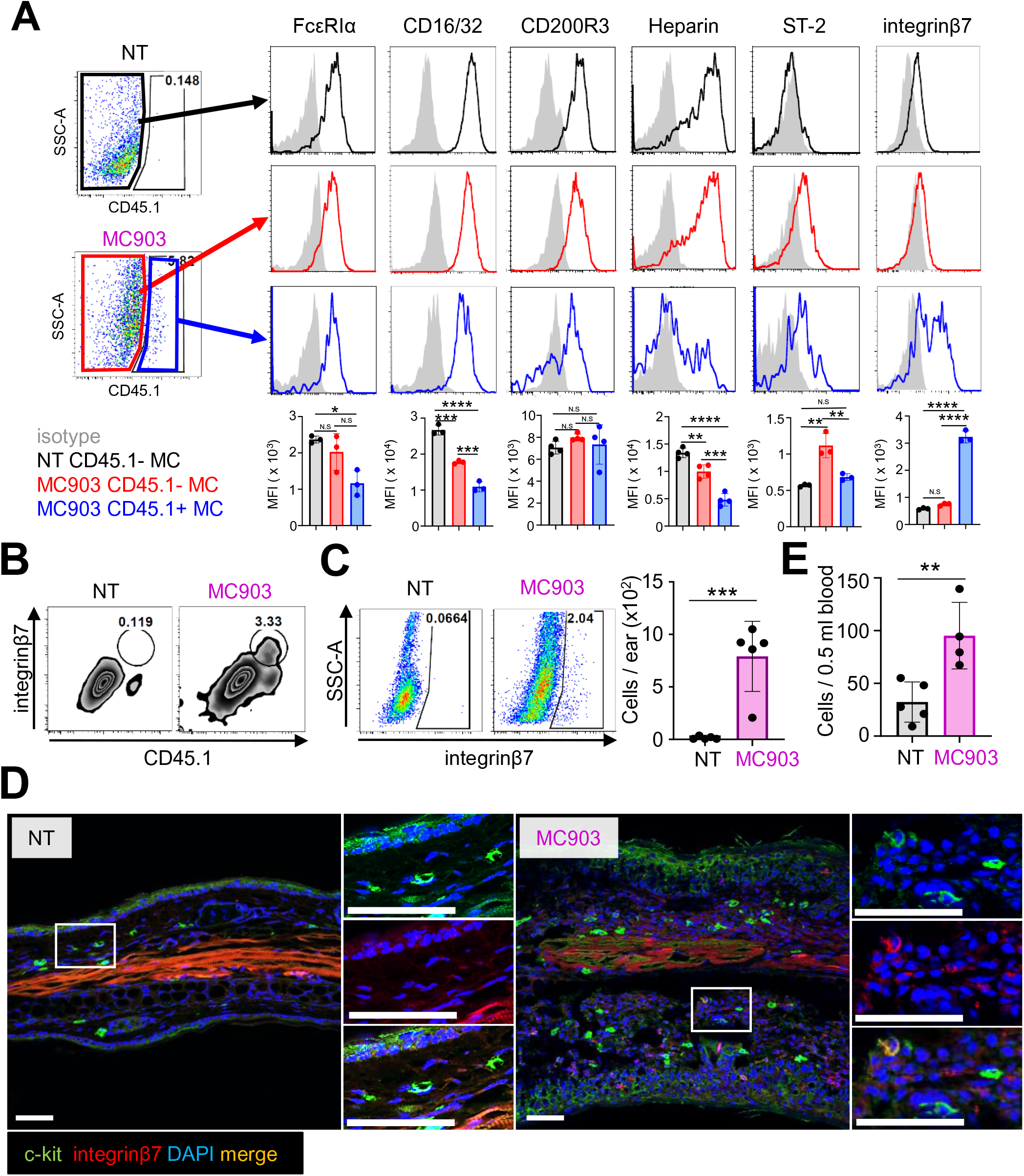
Identification of bone marrow (BM)-derived integrinβ7^+^ MCs in AD-like skin. **A,** Skin MCs are gated as Live^+^ CD45^+^ Lin^-^ CD11b^-^ CD11c^-^ c-kit^+^ cells. Representative flow cytometry plots and histograms for the expression of FcεRIα, CD16/32, CD200R3, heparin, ST-2, and integrinβ7 on CD45.1^-^ MCs in NT skin (black line) and CD45.1^-^ (red line) and CD45.1^+^ MCs (blue line) in MC903-trated skin. The gray area represents isotype control antibody. Mean fluorescent intensity (MFI) of each protein was shown as a bar graph (n = 3 or 4) **B,** Identification of integrinβ7^+^ MCs on CD45.1^+^ MCs in NT and MC903-treated skin. **C**, Identification and the number of integrinβ7^+^ MCs in NT and MC903-treated skin using WT mice. (n = 5) **D**, Immunostaining of c-kit (green), integrinβ7 (red), and DAPI (blue) in NT and MC903-treated skin. Scale bar, 50 μm. **E**, The number of MCps in 0.5 ml blood of NT and MC903-treated mice. (n = 4 or 5). Results are expressed as the mean ± SD. Data are representative of at least three independent experiments. *, p < 0.05, **, p < 0.01, ***, p < 0.001****, p < 0.0001, NS, not significant.

Integrinβ7^+^ MCs were all derived from BM cells (Figure 2B) and were detected only in AD-like skin (Figure 2C) using WT mice. Integrinβ7^+^ MCs were positive for FcεRIα and IgE, and partially negative for CD200R3 and heparin, which was consistent with the results of BM-derived MCs (Figure S2A). Immunohistochemistry also demonstrated c-kit^+^ integrinβ7^+^ cells in the deep dermis of AD-like skin (Figure 2D). Furthermore, although mature CTMCs (Lin^-^ CD11c^-^ CD11b^-^ CD200R3^+^ c-kit^+^ cells) were not detected in blood, the number of circulating MCps (Lin^-^ CD11c^-^ CD11b^-^ CD200R3^-^ c-kit^+^ integrinβ7 ^+^ CD16/32^+^ ST-2^+^ cells) (Figure S2B),(Dahlin et al., 2013) was significantly increased in mice treated with MC903 (Figure 2E). These data demonstrated that BM-derived MCs in AD-like skin are derived from circulating integrinβ7^+^ MCps, and integrinβ7 can be a marker to distinguish infiltrating BM-derived MCs from rMCs (integrinβ7^-^ MCs) in the skin. Integrinβ7^+^ MCs were also detected in the skin of other AD model (oxazolone-induced AD model), and psoriasis model (imiquimod-induced psoriasis model), but were not detected in DNFB-induced contact hypersensitivity model (Figure S2C).

### Integrinβ7^+^ MCs in AD-like skin showed transcript signatures of CTMCs and MMCs

To further characterize integrinβ7^+^ MCs (BM-derived MCps) in a comparison with rMCs (integrinβ7^-^ MCs), integrinβ7^+^ MCs and rMCS were isolated from NT and AD-like skin by flow cytometry sorting, then morphologically analyzed by Giemsa staining and electron microscopy. rMCs in NT skin contained compact basophilic granules in cyto-plasma (Figure 3A, left panels, Figure S3A, a left panel), which can be regarded as a resting state of MCs, while rMCs in AD-like skin were bigger and their cell surfaces became rough (Figure 3A, middle panels, Figure S3A, a middle panel), which might reflect their activating state.(Lorentz et al., 2012) On the other hand, integrinβ7^+^ MCs in AD-like skin contained less basophilic granules, suggesting the immaturity of the cells (Figure 3A, right panels, Figure S3A, a right panel). In addition, integrinβ7^+^ MCs did not show multi-segmented nucleus which are generally observed in basophils.

**Figure 3.**
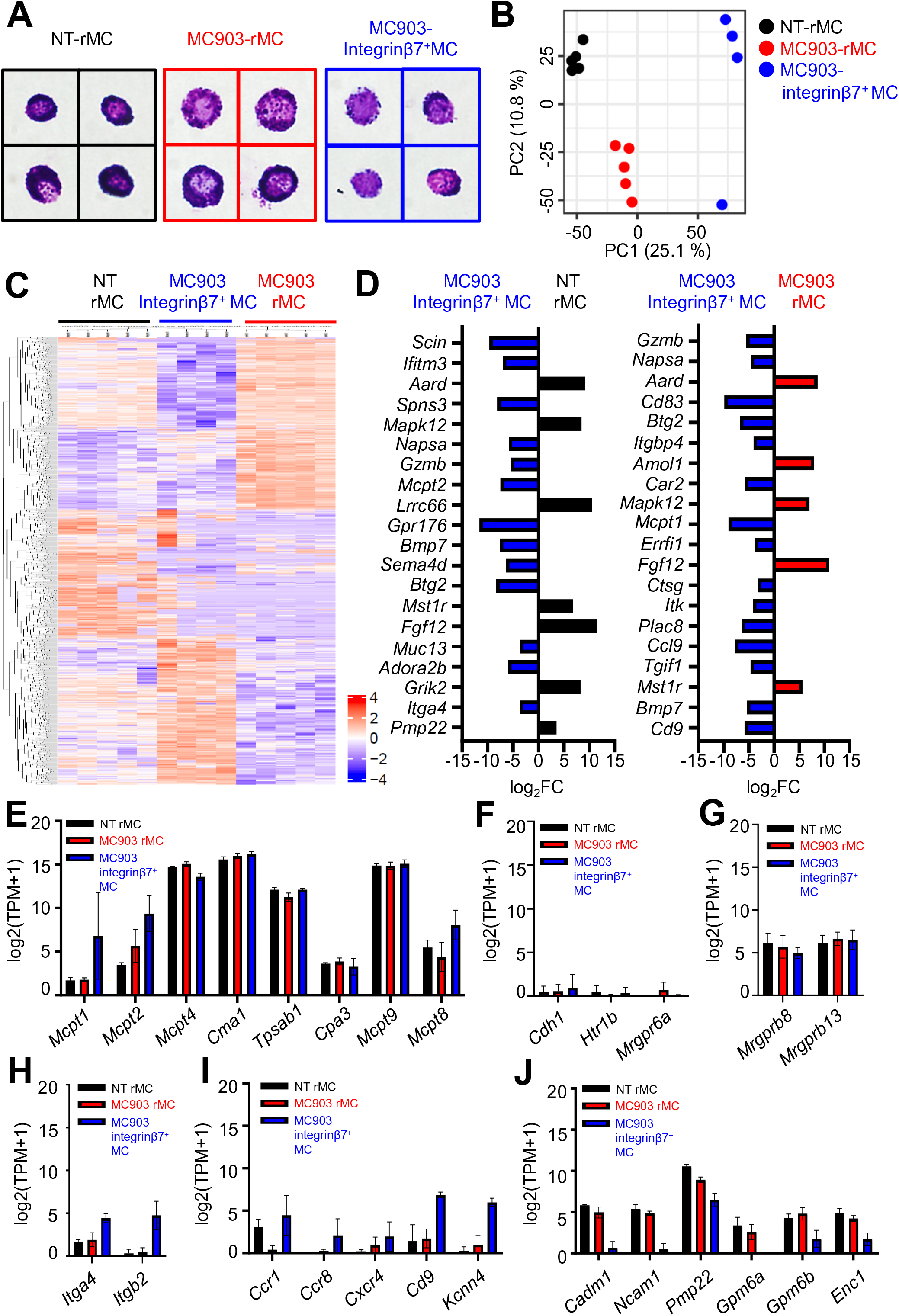
Integrinβ7^+^ MCs in AD-like skin showed signature transcripts of both MMCs and CTMCs. **A,** Giemsa staining (× 100) of rMCs and integrinβ7^+^ MCs in NT and MC903-treated skin. **B,** Principal component analysis of rMCs in NT skin (black dot), rMCs in MC903-treated skin (red dot), and integrinβ7^+^ MCs in MC903-treated skin (blue dot). Each colored circle represents sorted cells from each mouse. (n = 4 or 5) **C,** Heatmap of all DEGs between rMC in NT skin, rMC in MC903-treated skin, and integrinβ7^+^ MCs in MC903-treated skin. (FDR < 0.05, edgeR) **D,** Top 20 DEGs between rMC in NT skin and integrinβ7^+^ MCs in MC903-treated skin (left panel), and rMC in MC903-treated skin and integrinβ7^+^ MCs in MC903-treated skin (right panel). **E-J,** The expression of representative genes associated with MC proteases (**E**), basophils (**F**), skin CTMC-specific MRGPRS (**G**), leukocyte adhesion (**H**), MC migration (**I**), and neurons (**J**) as determined by RNA-seq are shown as bar graphs of log_2_ (TPM + 1) with the mean ± SD.

To investigate the difference in gene expression, RNA-seq was performed for each MC population. Principal component analysis (PCA) demonstrated three distinct MC populations (Figure 3B), and heatmap revealed 1157 differentially expressed genes (DEGs) (FDR > 0.05) (Figure 3C). Top 20 DEGs between integrinβ7^+^ MCs and rMCs in NT or AD-like skin highlighted several genes associated with MMC or MCp migration including *Mcpt1, Mcpt2* (specifically expressed in MMCs), *Itga4*, and *Cd9* (Figure 3D). Since MMCs and CTMCs contain different kinds of proteases (MMC; *Mcpt1, Mcpt2.* CTMC; *Mcpt4, Cma1, Tpsab1, Mcpt9*),(Gurish and Austen, 2012), (Derakhshan et al., 2021), (Pejler et al., 2010) we further compared the expression of these proteases among these MC populations. Integrinβ7^+^ MCs showed the increased expression of *Mcpt1*, and *Mcpt2*, while the expression levels of *Mcpt4*, *Cma1*, *Tpsab1*, and *Mcpt9* was almost the same as those of rMCs (Figure 3E). Integrinβ7^+^ MCs also expressed *Mcpt8*, a signature transcript of basophils, but the contamination of basophils is unlikely, because other basophil-specific transcripts, including *Cdh1*, *Htr1b*, and *Mrgpra6* were not expressed in integrinβ7^+^ MCs (Figure 3F).(Dwyer et al., 2017) Consistent with our results, the increased expression of *Mcpt8* was also reported in integrinβ7^+^ inducible MCs detected in the lung.(Derakhshan et al., 2021)

A previous report showed that CTMCs have several different transcripts dependent on peripheral tissues, and skin MCs specifically express *Mpgprb8* and *Mrgprb13* (Dwyer et al., 2017). Consistently, rMCs expressed *Mpgprb8* and *Mrgprb13,* and integrinβ7^+^ MCs also expressed them at similar levels (Figure 3G). In addition, integrinβ7^+^ MCs were enriched in genes related to MC migration, including *Itga4*, *Itgb2*, *Ccr1*, *Ccr8*, *Cxcr4*, *Cd9*, and *Kcnn4.* On the contrary, rMCs were increased in genes related to neurons including *Cadm-1*, *Ncam-1*, *Pmp22*, *Gpm6a*, *Gpm6b*, and *Enc* (Figure 3J), which may be consistent with a previous study showing the close localization of skin MCs to sensory nerves and contribution of Cadm-1 to their interactions with neurons.(Magadmi et al., 2019) Taken together, these data demonstrated that integrinβ7^+^ MCs showed both MMC signatures and skin type CTMC signatures, and they specifically expressed migration related genes.

We further compared gene expression of rMCs and integrinβ7^+^ MCs in NT and AD-like skin with progenitor cells in BM. A previous study demonstrated MCs and basophils are derived from E-cadherin^high^ granulocyte-monocyte progenitors (GMPs) in BM (Wanet et al., 2021). E-cadherin^high^ GMPs are further divided into two subsets by FcεRIα expression; pro-basophil and mast cell progenitors (BMPs) (E-cadherin^high^ FcεRIα^-^ GMPs) and pre-BMPs (E-cadherin^high^ FcεRIα^+^ GMPs).(Wanet et al., 2021) Using the public data (Wanet et al., 2021), we investigated the correlation of gene expressions in each MC population with pro-BMPs and pre-BMPs. Compared to rMCs in NT and AD-like skin, integrinβ7^+^ MCs were most correlated in gene expressions with both pro-BMPs and pre-BMPs (Figure S3B). This result supported our hypothesis that integrinβ7^+^ MCs in AD-like skin are derived from progenitor cells in BM.

### Circulating MCps migrated to allergic skin via LPAM-1

We next examined the adhesion molecules that mediate the recruitment of MCps into AD-like skin. RNA-seq data showed the increased expression of *Itga4* (encoding integrinα4) and *Itgb2(*encoding integrinβ2). The heterodimer of integrinα4 and integrinβ7 (LPAM-1) interacts with MadCAM-1 and/or VCAM-1 on blood endothelial cells and contribute to the migration of MCps to inflamed lung.(Berlin et al., 1995), (Abonia et al., 2006) Integrinβ2 also binds to other integrins, and the heterodimers are related to leukocyte adhesions,(M. Amin Arnaout, 1990) but its involvement in MCp migration remains unknown. First, the expression of integrinα4 and integrinβ2 on rMCs and integrinβ7^+^ MCs in NT and AD-like skin, and blood MCps were analyzed by flow cytometry. Integrinα4 was highly expressed on integrinβ7^+^ MCs in AD-like skin and blood MCps, and weakly expressed on rMCs (Figure 4A, upper panels). On the other hand, almost no expression of integrinβ2 were observed on rMC populations and blood MCps, and weak expression was observed on integrinβ7^+^ MCs (Figure 4A, lower panels). In addition, the expression of MadCAM-1 and VCAM-1, ligands of LPAM-1, was significantly increased on blood endothelial cells (Live^+^ CD45^-^ Ter119^-^ gp38^-^ CD31^+^ cells) in AD-like skin by flow cytometry (Figure 4B), suggesting the contribution of LPAM-1 to MCp migration to AD-like skin. To clarify this, blocking antibodies for LPAM-1 (anti-integrinα4β7 antibody) were systemically administered every other day from day 1 in the MC903- induced AD model. Although the ear swelling and the number of rMCs were not different in between isotype Ab- and blocking Ab-administered groups (Figure 4C), the number of integrinβ7^+^ MCs was significantly decreased in the blocking Ab-administered group (Figure 4D). These data demonstrated that the migration of circulating MCps to AD-like skin is mediated by LPAM-1 at least in part.

**Figure 4.**
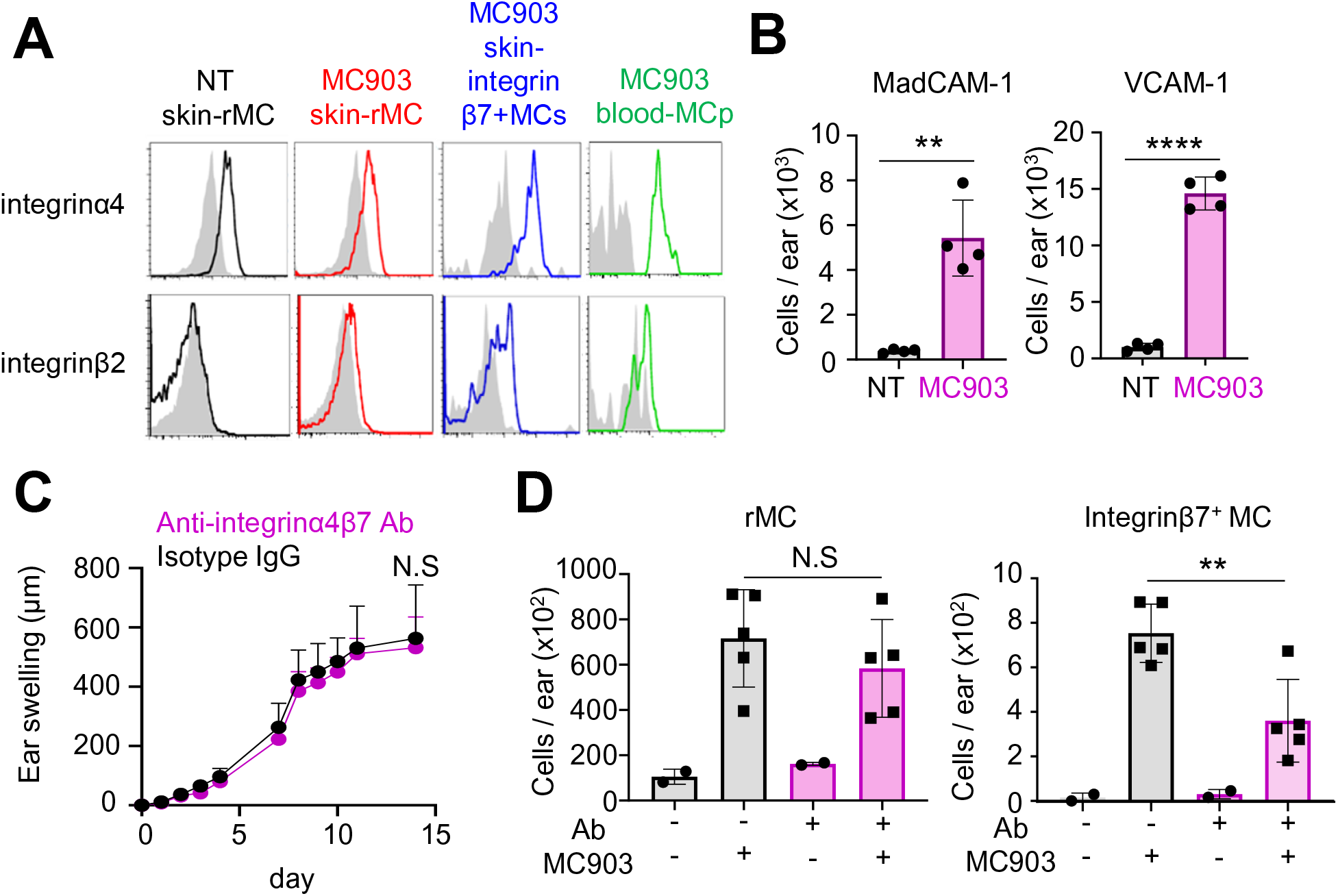
Circulating MCps migrate to AD-like skin via integrinα4β7. **A,** The expression of integrinα4 and integrinβ2 on rMCs in NT skin (black line), rMCs in MC903-treated skin (red line), integrinβ7^+^ MCs in MC903-treated skin (blue line), and on circulating MCps (green line). The gray area represents isotype control antibody. **B,** The numbers of MadCAM-1^+^ and VCAM-1^+^ blood endothelial cells (Live^+^ CD45^-^ Ter119^-^ gp38^-^ CD31^+^ cells) in NT and MC903-treated skin. (n = 4) **C,** Ear swelling of MC903-treated skin with/without blocking antibody against LPAM-1. (n = 2 for NT and n = 5 for MC903-treated skin) **D,** The number of rMCs and integrinβ7^+^ MCs in NT and MC903-treated skin with/without antibody treatment. Anti-integrinα4β7 mAb (300 μg/body) or isotype IgG (300 μg/body) were administered intra-peritoneal every other day from day 1. (n = 2 for NT and n = 5 for MC903) Results are expressed as the mean ± SD. Data are representative of at least three independent experiments. **, p < 0.01, ***, p < 0.001, and ****, p < 0.0001. NS, not significant.

### Integrinβ7^+^ MCs are dispensable for MC903-induced AD-like skin inflammation

To analyze function of integrinβ7^+^ MCs in MC903-induced AD-like skin, we first compared the gene expression of representative cytokines and chemokines with RNA-seq data. Integrinβ7^+^ MCs demonstrated the slightly increased expression of *Il10*, while the expression levels of *Tnf, Il4, Il2, and Tgfb* were almost the same as those of rMCs (Figure 5A). However, flow cytometry analysis showed IL-10 was slightly expressed in rMCs in NT skin, while IL-10 was not detected in rMCs or integrinβ7^+^ MCs in AD-like skin (Figure S4A). Integrinβ7^+^ MCs also showed the increased expression of *Ccl3, Ccl4, and Ccl9,* while rMCs showed the increased expression of *Ccl7* (Figure 5B). These data suggested that integrinβ7^+^ MCs might have different immunological functions from rMCs. Next, to evaluate *in vivo* function of integrinβ7^+^ MCs in AD-like skin, we first compared the ear swelling between Wsh/Wsh mice, which lack both resident and integrinβ7^+^ MCs (BM-derived MCs), and WT mice. Wsh/Wsh mice and WT mice were co-housed more than two weeks before MC903 treatment for skin microbiome to be shared between each mouse strain.(Kobayashi et al., 2019) As a result, ear swelling was slightly attenuated in Wsh/Wsh mice, but there was no significant difference in the number of eosinophils, one of the key cells of MC903-induced AD-model (Figure 5E).(Naidoo et al., 2018) This result suggested that either or both rMCs and integrinβ7+ MCs have a pro-inflammatory role irrespective of eosinophils in MC903-induced AD model. We next compared the ear swelling between Mas-TRECK mice and littermate-WT mice. Using Mas-TRECK mice in which MCs and basophils can be depleted by diphtheria toxin (DT), we intradermally (i.d.) injected DT to deplete only resident MCs in the ear skin, and then applied the mice to the MC903-induced AD like model (Figure S4B). Similar to experiments using Wsh/Wsh mice, ear swelling showed a tendency of the slight decrease in Mas-TRECK mice with i.d. DT, but there was no significant difference in the number of eosinophils (Figure 5F). This result suggested that rMCs promote the inflammation, while integrinβ7+ MCs do not compensate for their function. Lastly, we generated BM-chimera mice in which BM cells from either Wsh/Wsh mice or WT mice are intravenously injected to lethally irradiated WT mice: Wsh/Wsh > WT or WT > WT (Figure S4C). Since skin resident MCs are radio-resistant, only integrinβ7^+^ MCs (BM-derived MCs) are deficient in Wsh/Wsh > WT mice, which enable us to analyze the functions of integrinβ7^+^ MCs (BM-derived MCs). Unlike Wsh/Wsh mice or Mas-TRECK mice with i.d DT, any difference in the ear swelling or the number of eosinophils was observed (Figure 5G). This result suggested that integrinβ7+ MCs are unnecessary for the AD-like inflammation and rMCs cover the function of integrinβ7+ MCs. Furthermore, considering that integrinβ7^+^ MCs infiltrate to AD-like skin on day 14, we examined the ear swelling responses at the remission phase of inflammation (after discontinuation of MC903 application) and skin fibrosis by measuring hydroxyproline. However, we did not see any difference in these parameters as well (Figure S4D). Taken together, rMCs showed a slight pro-inflammatory role on MC903-induced AD-like skin, while integrinβ7^+^ MCs did not show an apparent role.

**Figure 5.**
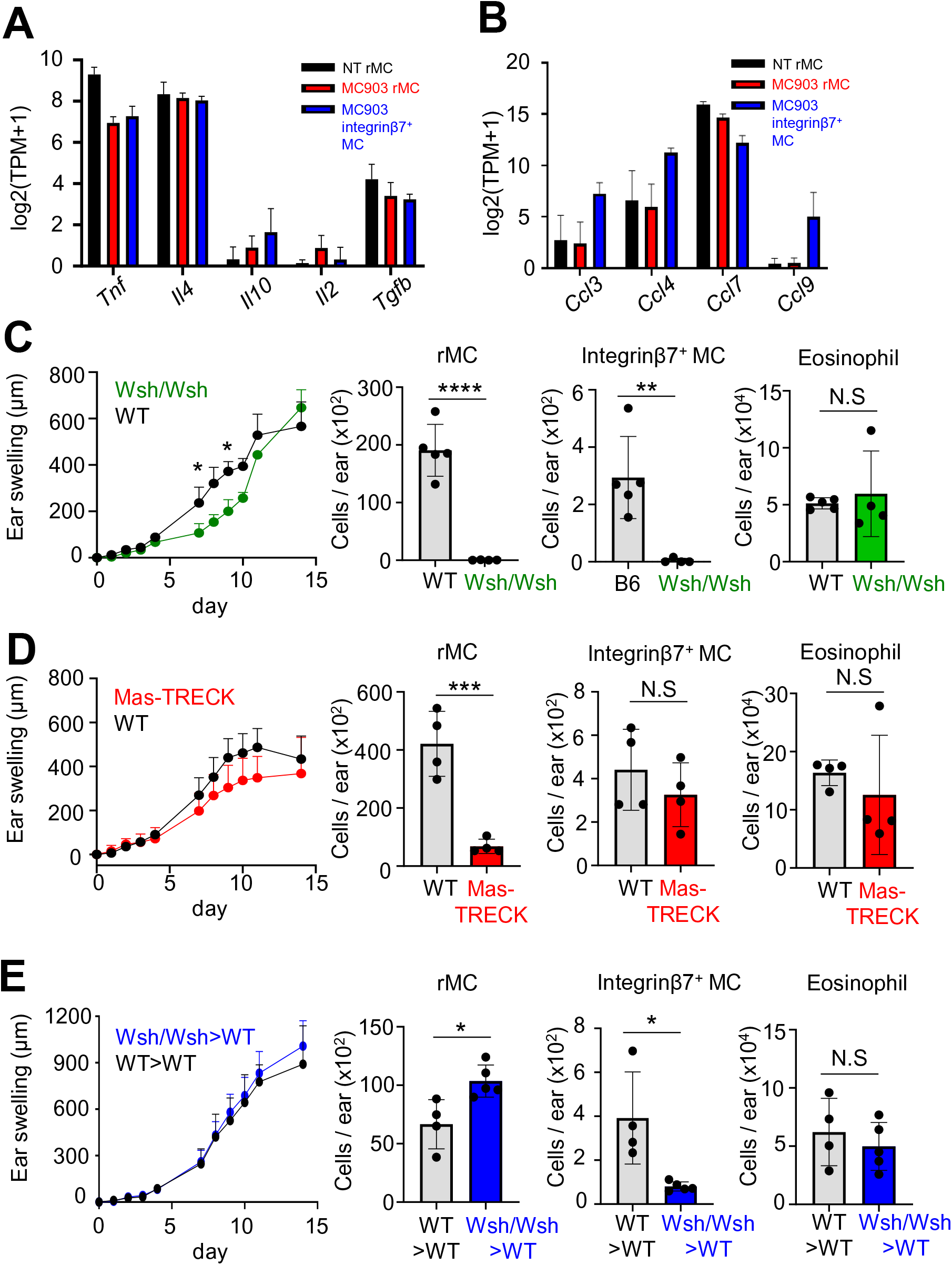
Integrinβ7^+^ MCs are dispensable for MC903-induced AD-like skin inflammation. **A, B,** The expression of representative genes of cytokines (**A**) and chemokines (**B**) as determined by RNA-seq are shown as bar graphs of log2 (TPM + 1) with the mean ± SD. **C,** Ear swelling responses and the number of rMCs, integrin β7^+^ MCs, and eosinophils in the skin on day14 in WT and Wsh/Wsh mice in MC903-induced AD model. (n= 4 or 5) **D,** Ear swelling responses and the number of rMC, integrin β7^+^ MCs, and eosinophils in the skin on day14 in WT and MasTRECK mice with i.d DT in MC903-induced AD model. (n= 4) **E,** Ear swelling responses and the number of rMC, integrin β7^+^ MCs, and eosinophils in the skin on day14 in WT>WT and Wsh/Wsh>WT mice in MC903-induced AD model. (n= 4 - 5) Results are expressed as the mean ± SD. Data are representative of at least two independent experiments. *, p < 0.05, **, p < 0.01, ***, p < 0.001****, p < 0.0001, NS, not significant.

### Integrinβ7^+^ MCs proliferated in AD-like skin and acquired CTMC phenotype

To investigate the fate of integrinβ7^+^ MCs after migration to AD-like skin, the number of BM-derived MCs (CD45.1^+^ MCs) and their cell surface markers were tracked on day 14, 21, 28, and 35 in the MC903-induced AD model by using CD45.1 chimera mice (Figure 6A). MC903 was applied to the ear skin for 10 days (Figure 1A). Ear thickness gradually decreased after day 14 and went back to almost normal levels on day 35 (Figure 6B). While the number of CD45.1^-^ MCs (rMCs) increased only until the peak of inflammation (day 14) (Figure 6C), the number of CD45.1^+^ MCs (BM-derived MCs) constantly increased over time even after the peak of inflammation (Figure 6D). On the other hand, the number of integrinβ7^+^ MCs in CD45.1^+^ MCs increased from day 0 to day 21, accounting for about 50% of CD45.1^+^ MCs, and then decreased on day 28 (Figure 6E). These data suggest that the migration of MCps to AD-like skin occurred mainly during from day 0 to day 21, and then the infiltrating MCs proliferated and stayed in the skin losing their integrinβ7 expression. Consistently, integrinβ7^+^ MCs showed an increased percentage of Ki67 positive populations compared to rMCs (Figure S5A). It is of note that basophils (Live^+^ CD45^+^ CD11b^-^ CD200R3^+^ c-kit^-^ cells) were almost absent in NT skin (day 0), and increased until day 14 then decreased afterwards. Basophils were mostly derived from BM (average 98.1 % on day 14) (Figure S5B). This demonstrate skin MCs and basophils are differently maintained in AD-like skin.

**Figure 6.**
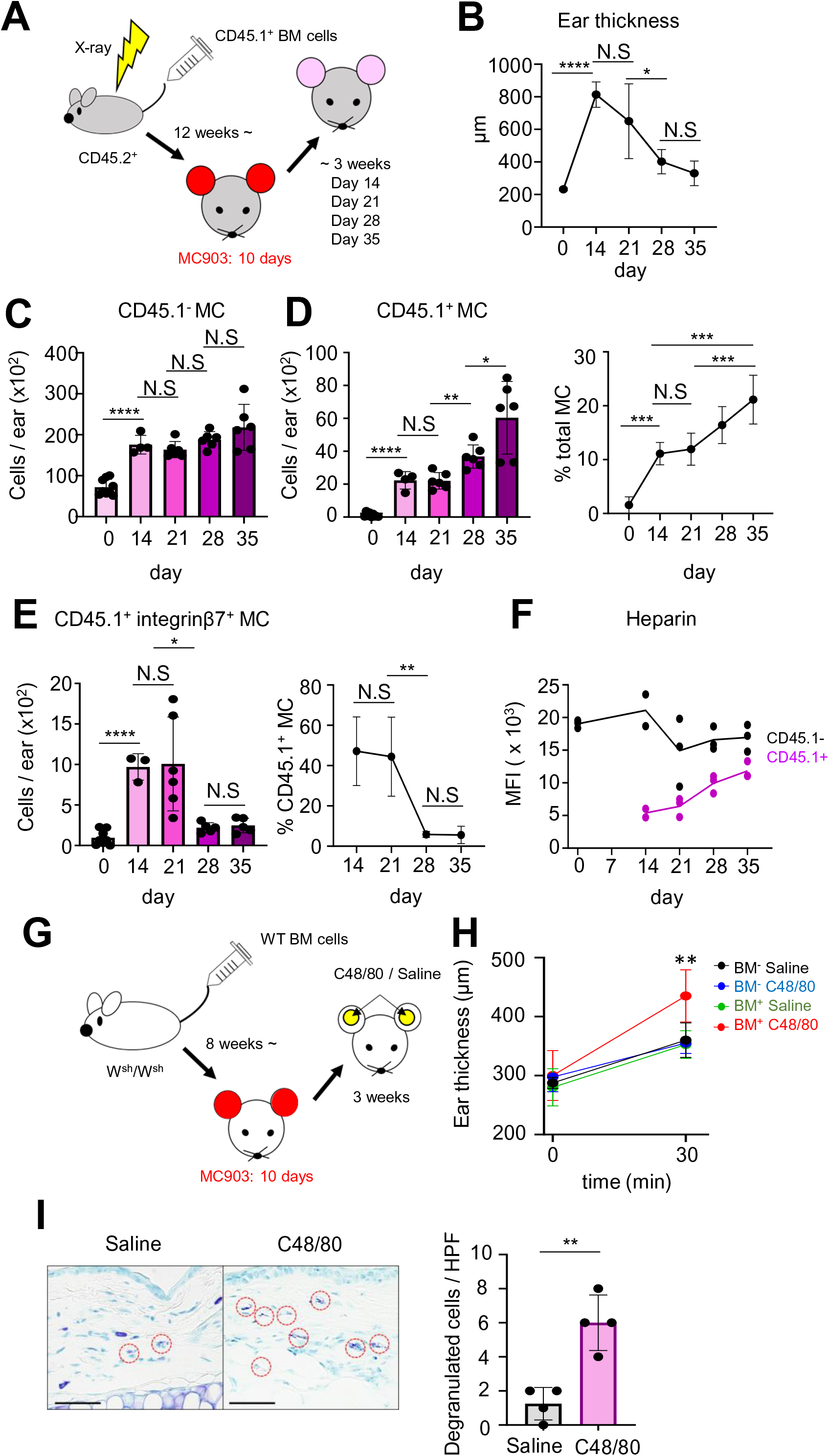
Integrinβ7^+^ MCs proliferated and differentiated into CTMCs in AD-like skin at the resolution phase of the inflammation. **A,** The experimental scheme of BM-chimera mice. **B,** Ear thickness on day 0, 14, 21, 28, and 35 in MC903-induced AD model. (n=4 - 6) **C,** The numbers of CD45.1^-^ MCs in the skin on day 0, 14, 21, 28, and 35 in MC903-induced AD model. (n = 4 - 6) **D,** The number and the ratio of CD45.1^+^ MCs to all MCs in the skin on day 0, 14, 21, 28, and 35. (n = 4 - 6) **E,** The number and the ratio of integrinβ7^+^ MCs to CD45.1^+^ MCs in the skin on day 0, 14, 21, 28, and 35. (n = 3 - 6) **F,** The MFI of heparin in CD45.1^+^ MCs in skin on day 0, 14, 21, 28 and 35. (n = 2 or 3) **G,** The experimental scheme of adoptive transfer of BM cells to Wsh/Wsh mice. **H,** Ear thickness before and 30 min after intradermal (i.d.) injection of saline or C48/80 in mice with or without BM transfer. **I,** Toluidine blue staining of the ear skin treated with saline or C48/80. The degranulated MCs are shown with red dotted circles. Scale bar, 50 μm. The numbers of degranulated MCs per high-power field (HPF) (× 40) (right panel) (n = 4). Results are expressed as the mean ± SD. Data are representative of at least two independent experiments. *, p < 0.05, **, p < 0.01, ***, p < 0.001, ****, p < 0.0001. NS, not significant.

To clarify whether integrinβ7^+^ MCs differentiate to CTMCs in AD-like skin, the expression of heparin was analyzed, because not MMCs but CTMCs express heparin. The expression level of heparin on CD45.1^+^ MCs were significantly increased from day 14 to day 35 (Figure 6F), suggesting maturation of infiltrating MCps to CTMCs. To investigate whether infiltrating MCps acquire CTMC phenotype in functions, the reaction against compound 48/80 (C48/80) was assessed, because degranulation in response to C48/80 is generally observed only in CTMCs.(Stanovnik et al., 1988) First, MC-deficient Wsh/Wsh mice were adoptively transferred with 1×10^7^ WT-BM cells to reconstitute BM-derived MCs. MC903 was applied to both ears for 10 days 8 weeks after the adoptive transfer. As we observed in CD45.1 BM-chimera mice, skin MCs were significantly increased only in AD-like skin in mice with BM-cell transfer (Figure S5C). Whole-mount immunostaining of ear skin showed c-kit^+^ cells gathered in some places of the ears, but were mainly observed around the bottom to the center of the ears (Figure S5D). C48/80 or saline was i.d. injected in those ears, and evaluation of ear swellings and histological analysis of MC degranulation were performed (Figure 6G). In BM-transferred mice, the ear swelling was prominent in ears injected with C48/80 than ears injected with saline (Figure 6H). Histological analysis also showed the increased number of degranulated MCs in C48/80-treated ears of BM-transferred mice (Figure 6I). Taken together, these data demonstrate that integrinβ7^+^ MCs can acquire CTMC phenotypes and elicit their physiological functions in the skin.

### c-kit^+^ FcεRI^+^ integrinβ7^+^ cells were observed in human AD skin

Finally, we examined whether infiltrating MCs can be detected in human AD skin. Human MCps are identified as Lin^-^ CD34^hi^ CD117^int/hi^ FcεRI^+^ cells in blood and express integrinβ7.(Dahlin et al., 2016) Therefore, the expression of c-kit, FcεRI, and integrinβ7 in human AD skin was analyzed by immunohistochemistry. In normal skin, c-kit^+^ FcεRI^+^ integrinβ7^+^ cells were not detected, while there were c-kit^+^ FcεRI^+^ integrinβ7^+^ cells in the reticular layer of the dermis of AD skin (Figure 7A). The number of dermal c-kit^+^ FcεRI^+^ cells was significantly higher in AD skin (Figure 7B), and an average of 4.2% of dermal c-kit^+^ FcεRI^+^ cells was positive for integrinβ7 in AD skin (Figure 7C). These data suggest that circulating MCps infiltrate to human AD skin as well.

**Figure 7.**
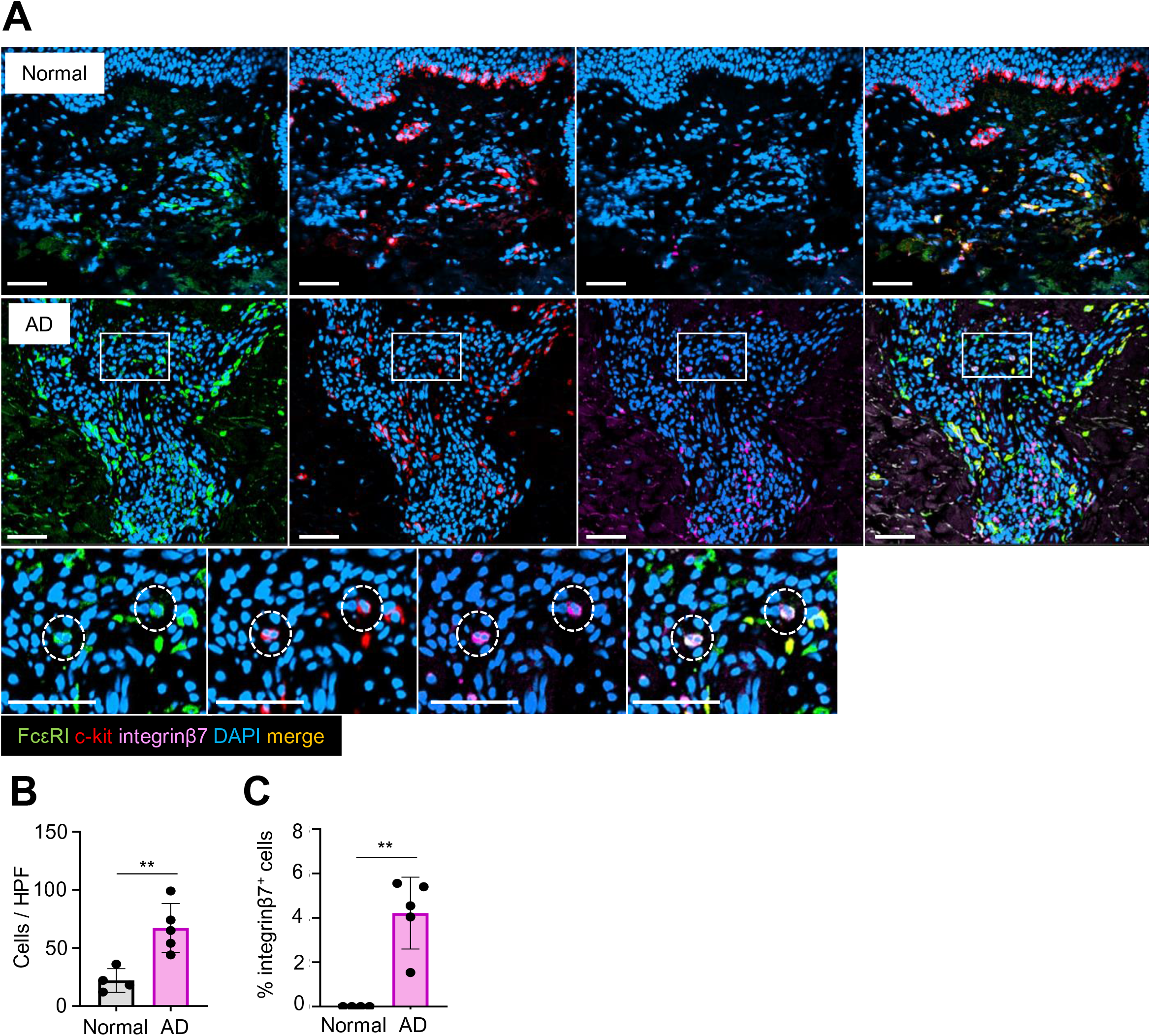
The expression of c-kit, FcεRI, and integrinβ7 in normal and AD skin. **A,** Immunostaining of FcεRI (green), c-kit (red), integrinβ7 (magenda), and DAPI (blue) in the human adult normal and AD skin. Triple-positive cells are shown with white dotted circles. Scale bar, 50 μm. **B,** The number of c-kit^+^ FcεRI^+^ cells in the dermis per HPF (× 20). **C,** The ratio of c-kit/FcεRI/integrinβ7-triple-positive cells among c-kit/FcεRI-double-positive cells in the dermis. (n = 4 for normal skin and n = 5 for AD skin) Results are expressed as the mean ± SD. **, p < 0.01.

## Discussion

Here, we revealed how skin MCs are maintained under allergic inflammation. Skin MCs were increased in AD-like skin due to local proliferation of rMCs and infiltration/proliferation of BM-derived circulating MCs. Infiltrating MCs were identified as integrinβ7^+^ MCps and showed an immature phenotype of CTMCs. Integrinβ7^+^ MCs proliferated and acquired the phenotype of CTMC in the skin.

The origin of skin MCs has been believed to be BM-derived HSCs, because skin-type CTMCs are inducible from BM cells *in vitro*,(Takano et al., 2008) and i.d. injection of BMMCs results in reconstitution of skin MCs in MC-deficient Wsh/Wsh mice.(Wolters et al., 2005) However, this concept has also been questioned, because intravenous transfer of BMMCs to Wsh/Wsh results in minimal reconstitution of skin MCs.(Kitamura et al., 1978) This was assumed that BMMCs and/or skin lack the expression of molecules essential for the migration, and skin MCs are maintained independently from BM supply. Then, the recent study by Gentek et al. revealed that adult skin MCs are derived from aorta-gonads-mesonephros region-derived HSCs, and are maintained locally by self-proliferation without migration of BM-derived MCps.(Gentek et al., 2018) However, it had been unanswered whether skin MCs are maintained only by local proliferation in inflammatory state such as AD. Our current study revealed that skin MCs are maintained not only by local proliferation but also supply from circulating MCps, further advancing our understanding of MC physiology. Increased expression of adhesion molecules, such as MadCAM-1 and/or VCAM-1, on blood endothelial cells seemed to partially mediate the infiltration of circulating MCps into skin, because blocking antibody for integrins decreased the number of MCps in the skin. Although we did not perform detailed analysis of chemotactic factors of MCps, some lipid mediators including PGE2 and LTB4,(Weller et al., 2007) or chemokine receptors as indicated by RNA-seq analysis, such as CCR1, CCR8, CXCR4, CD9, or KCNN4, may be possible candidates as those factors.

Infiltration of MCps into the tissue under an inflammatory state and their expression of integrinβ7 has recently been reported in an allergic inflammation model in the lung.(Abonia et al., 2006) Their gene transcripts showed MMC-related signatures which are completely distinct from CTMCs.(Derakhshan et al., 2021) Similar to their results, integrinβ7^+^ MCs in AD-like skin showed increased expression of genes related to MMCs including *Mcpt1* and *Mcpt2*. However, integrinβ7^+^ MCs in the AD-like skin also expressed skin specific CTMC signature transcripts including *Cd200r3*, *Mcpt4*, *Mrgprb18*, *and Mrgprb13*. This supports the conventional notions that MCps have the potential to differentiate into both MMCs and CTMCs, and terminal differentiations are dependent on the microenvironment where they migrate.(Metcalfe, 1997)

There are some limitations of our study. First, although we found two MC populations in AD-like skin (rMCs and integrinβ7^+^ MCs), their functional differences were not clarified. It was already known that skin MCs are increased in AD skin, and both pro- and anti-inflammatory functions had been reported.(Kawakami et al., 2009) For instance, skin MCs facilitate house dust mite-induced allergic inflammation by interaction with peripheral neurons.(Serhan et al., 2019) On the other hand, MCs suppress hapten-induced chronic allergic dermatitis by activating regulatory T cells via IL-2 production in spleen.(Hershko et al., 2011) These functional differences could be explained due to the different experimental models and/or locations where MCs function. In MC903-induced AD-like model, rMCs showed a slight pro-inflammatory role, while integrinβ7^+^ MCs did not show a clear role. This might be because the number of integrinβ7^+^ MCs is quite small in the skin. Considering that integrinβ7^+^ MCs eventually differentiate into skin type CTMCs, the function of integrinβ7^+^ MCs might be subsidiary to rMCs. Although we cannot exclude the possibilities that BM-derived MCs have specific functions in other dermatitis models, further analysis about their *in vivo* functions will be our future scope. Second, how MCps adapt to skin, including proliferation and differentiation, was not fully addressed. Signaling from SCF is a possible mechanism of the proliferation/differentiation of MCps in our model, because SCF is crucial for CTMC differentiation and proliferation, and is expressed by various cells in the skin including keratinocytes.(Wang et al., 2017) In fact, the expression of SCF was increased in the AD in our model. In addition to SCF, interaction with fibroblasts may also be important for MCp differentiation and proliferation, because in vitro studies demonstrated that skin-type CTMCs are inducible from BM cells when cocultured with fibroblasts and SCF.(Takano et al., 2008) Although the precise mechanism of how fibroblasts induce the differentiation is not known, CD44, a primary receptor for hyaluronan, is reported to be involved in MC proliferation in the skin.(Takano et al., 2009) These points are also open questions to be answered.

In conclusion, our study revealed that skin MCs are maintained by the proliferation of rMCs, and migration of circulating MCps into the AD-like skin and their local adaptability. These findings could give a novel insight into the understanding of the heterogeneity of MCs in various inflammatory skin diseases.

## Materials and Methods

### Mice

All experiments were performed using 6 to 12-week-old C57BL/6-background female mice (OrientalBioService, Kyoto, Japan). C57BL/6-*KitW_-sh_/W_-sh_* (Wsh/Wsh) mice were provided by RIKEN BioResource Research Center (Ibaraki, Japan). Mas-TRECK mice were generated and kindly provided by Dr. Masato Kubo (RIKEN Yokohama Institute).(Otsuka et al., 2011), (Sawaguchi et al., 2012) All mice were maintained under specific pathogen-free conditions in the Institute of Laboratory Animals at Kyoto University Graduate School of Medicine. All experimental procedures were approved by the Institutional Animal Care and Use Committee of the Kyoto University Graduate School of Medicine.

### MC903-induced murine atopic dermatitis (AD) model

MC903, calcipotriol (R&D Systems, Minneapolis, MN), was used to induce AD-like skin in mice. Both sides of each ear were topically applied with 2 nmol MC903 in 20 μl ethanol for 10 days with 2 days break after 5 consecutive days. Ear thickness was measured every day using a thickness gauge, and the difference from day 0 was expressed as the change in ear thickness.

### Preparation of single-cell suspensions

The dorsal halves of the ear skin of mice were collected and incubated for 70 min at 37°C in cRPMI containing 0.33 mg/ml Liberase TL (Roche, Basel, Switzerland) and 0.05% DNase I (Sigma-Aldrich, St. Louis, MO) dissolved in RPMI 1640 (Invitrogen, Carlsbad, CA) containing 10% FCS. Digested skin sheets were homogenized and filtered using a 40 µm cell strainer (BD Biosciences, San Jose, CA) to obtain single-cell suspensions. 500 μl of peripheral blood was collected from each mouse by puncturing the submandibular vein with an 18G needle into EDTA-treated tubes. Red blood cells were lysed by using a lysis buffer (0.15 M NH_4_Cl; 14.3 mM NaHCO_3_; 0.1 mM Na_2_EDTA; pH 7.3).

### Flow cytometry

Nonspecific antibody binding was blocked with an anti-CD16/32 antibody (2.4G2; BD Biosciences, San Jose, CA). Cells were stained with the following antibodies: anti-mouse antibody to CD45 (30-F11; BD Biosciences), CD45.1 (A20; BioLegend, San Diego, CA), CD3e (145-2C11; Thermo Fisher Scientific, Waltham, MA), B220 (RA3-6B2, BD Biosciences), Ter119 (TER-119; BD Bioscience), Siglec-F (E50-2440; BD Biosciences), Ly-6G (1A8; BD Biosciences), CD11b (M1/70; BD Biosciences), CD11c (N418; BioLegend), CD200R3 (Ba13; BioLegend), c-kit (2B8; BD Biosciences), FcɛRIα (MAR-1; eBioscience, San Diego, CA), ST-2 (DJ8; MyBioSource, San Diego, CA), integrinβ7 (FOB504; BD Biosciences), integrinα4 (R1-2; Invitrogen), integrinβ2 (M18/2; BioLegend), IgE (R35-72; BD Biosciences), CD31 (MEC13.3; BioLegend), gp38 (8.1.1; BioLegend), VCAM-1 (429; BioLegend), MadCAM-1 (MECA-367; BioLegend), Ki67 (SolA15; eBioscience), IL-10 (JES5-16E3; BD Biosciences). Fixable viability dye eFluor 780 (FVD) (Thermo Fisher Scientific) was used to gate out dead cells. Skin MCs were gated as FVD^-^ CD45^+^ Lin (CD3, B220, Ter119, Siglec-F, Ly-6G)^-^ CD11b^-^ CD11c^-^ c-kit^+^ cells. Blood MCps were gated as FVD^-^ CD45^+^ Lin^-^ CD11b^-^ CD11c^-^ CD200R3^-^ c-kit^+^ ST- 2^+^ CD16/32^+^ cells. Skin blood endothelial cells were gated as FVD^-^ (Live^+^) CD45^-^ Ter119^-^ gp38^-^ CD31^+^ cells. Skin basophils were gated as FVD^-^ CD45^+^ CD11b^-^ CD200R3^+^ c-kit^-^ cells. Flow cytometry was performed using an LSRFortessa cell analyzer (BD Biosciences) and analyzed with FlowJo (TreeStar, San Carlos, CA). Cell numbers of each cell subset were evaluated by using Flow-Count Fluorospheres (Beckman Coulter, Brea, CA) and presented as numbers per tissue after exclusion of doublet and dead cells.

### Whole-mount staining of the murine ear skin

The dorsal halves of the ear skin were collected and floated for 30 minutes at 37°C on 35.56 mg/ml ammonium thiocyanate (Wako, Osaka, Japan) dissolved in PBS to separate the epidermis and dermis. Then, the dermis was fixed with 4% paraformaldehyde for 1 hour at RT, washed with PBS, and stained overnight at 4°C with avidin-FITC (BioLegend), anti-human/mouse CD117 (c-kit) antibody (R&D Systems Minneapolis, MN) and anti-CD31 antibody (MEC13.3; BioLegend). Then samples were stained with Alexa Fluor 594 anti-goat IgG antibody (Invitrogen) for 1 hour at room temperature. Samples were mounted with antifade mounting medium (Thermo Fisher Scientific). Fluorescent images were analyzed with a Nikon confocal microscope (Nikon A1RMP, Nikon, Tokyo, Japan).

### CD45.1-BM chimeric mice

Recipient CD45.2^+^ wild type (WT) mice were lethally irradiated with 9.5 Gy by an X ray irradiation device and received 5×10^6^ BM cells from congenic CD45.1^+^ mice intravenously through the tail vein. The chimeric mice were subjected to the indicated experiments over 12 weeks after the reconstitution.

### Parabiosis

Pairs of parabiotic mice consisting of CD45.2^+^ WT and congenic CD45.1^+^ mice were generated as described previously.(Duyverman et al., 2012) The parabiotic pairs were treated with MC903 two weeks after the surgery. Percentages of CD45.1^+^ and CD45. 2^+^ cells among blood and skin neutrophils were analyzed in each mouse to confirm efficient blood mixing.

### Enzyme-linked immunosorbent assay (ELISA) for the detection of skin SCF

All protein was extracted from ear skin samples by homogenizing tissues in RIPA buffer containing protease inhibitor cocktails (Sigma-Aldrich, St. Louis, MO). SCF levels were determined using ELISA kit (R&D Systems). All procedures were performed according to the manufacturer’s protocol. The absorbance at 450 nm was measured with a microplate reader (FlexStation 3; Molecular Devices, San Jose, CA).

### Immunostaining of the murine ear skin

Mouse ears were fixed in 4% paraformaldehyde (PFA), and embedded in O.C.T. compound (Sakura Finetek, Torrance, CA). Five micrometer slices were treated with Image-iT FX signal enhancer (Invitrogen), and incubated with anti-human/mouse CD117 (c-kit) antibody (R&D Systems), anti-integrinβ7 antibody (FIB504; BioLegend) for overnight at 4℃. Then samples were stained with Alexa Fluor 488 anti-goat IgG antibody (Invitrogen) and Alexa Fluor anti-rat IgG antibody (Invitrogen) for 1 hour at room temperature. The slices were mounted with ProLong Diamond Antifade Mountant with 4’,6-diamino-2-phenylindole (DAPI) (ThermoFisher Scientific). Fluorescent images were analyzed with a Nikon confocal microscope (Nikon A1RMP, Nikon).

### Oxazolone-induced chronic allergic dermatitis model

Oxazolone (Wako) was used as a hapten. Mice were sensitized with 25 µl of 3% (wt/vol) oxazolone in ethanol on their shaved abdomens. Five days after sensitization, 20 µl of 0.6% (wt/vol) oxazolone was applied to both sides of ears three days in a week total of 10 times. Ear thickness was measured for each mouse before and 24 hours after the last hapten application using a thickness gauge.

### Imiquimod-induced psoriasis-like dermatitis model

10 mg of imiquimod-containing cream, Aldara (Beselna Cream 5%, Mochida Pharmaceuticals, Tokyo, Japan), was applied onto the mouse ear for 7 consecutive days. Ear thickness was measured for each mouse every day using a thickness gauge, and the difference from day 0 was expressed as the change in ear thickness.

### DNFB-induced acute contact hypersensitivity model

1-Fluoro-2,4-dinitrobenzene (DNFB) (Nacalai Tesque, Kyoto, Japan) was used as a hapten. Mice were sensitized with 25 µl of 0.5% (wt/vol) DNFB in 4:1 acetone/olive oil (vol/vol) on their shaved abdomens, and then elicited with 20 µl of 0.2% (wt/vol) DNFB on both sides of each ear 5 days after sensitization. Ear thickness was measured for each mouse before and 24 hours after elicitation using a thickness gauge.

### MC isolation from skin

CD45^+^ cells from ear skin were enriched by CD45 MicroBeads (Miltenyi Biotec, Bergisch Gladbach, Germany) according to the manufacturer’s protocol. Then, rMCs and integrinβ7^+^ MCs were sorted using FACS AriaII cell sorter (BD Biosciences).

### Giemsa staining

Sorted MC cell suspensions were stained with Giemsa Stain (Sigma-Aldrich) following the standard protocol and examined by light microscopy.

### Electron microscopy

Sorted MC cells suspensions were resuspended with iPGEL (NIPPON Genetics, Tokyo, Japan), then were prefixed with half Karnovsky solution containing 2% PFA and 2% glutaraldehyde (Wako) in 0.1 M cacodylic acid (Wako) overnight at 4 °C and fixed in 1% osmium before the dehydration and epon embedding. Samples were then ultrathin-sliced and observed using transmission electron microscope (H-7650, Hitachi High-Technologies Corporation, Tokyo, Japan).

### In vivo blocking of integrinα4β7

Anti-integrinα4β7 (LPAM-1) Ab (DATK32) and rat IgG2a isotype control (2A3) were purchased from Bio X Cell (West Lebanon, NH). Mice were injected intraperitoneally with 300 μg of anti-LPAM-1 Ab or rat IgG2a isotype control every other day from day 1 a total of 7 times.

### Histological examination of the murine ear skin

Skin tissues were fixed with 10% formalin in PBS and embedded in paraffin. Sections with a thickness of 5 μm were prepared, stained with toluidine blue, and examined by light microscopy.

### RNA-seq and data analysis

Up to 100 cells of resident mast cells (rMCs) and integrinβ7^+^ MCs from both ears of NT mice (n = 5) and MC903-treated mice (n = 6) were collected into CDS sorting solution (Takara Bio Inc, Shiga, Japan) using 96 well plate. Then cDNA was synthesized using SMART-Seq HT (Takara Bio Inc) protocol, and purified using Agencourt AMPure XP (Beckman Coulter). RNA-seq libraries were prepared using Illumina Nextera XT DNA Library Prep Kit (Illumina, San Diego, CA) and sequencing was performed on HiSeq 4000 platform (Illumina). Sequencing data in the form of FASTQ files were mapped onto the Mus musculus genome build GRCm38 using the STAR aligner (version 2.5.2b) with gene annotations obtained from GENCODE (version M22). One sample from rMCs of MC903-treated skin and two samples from integrinβ7^+^ MCs of MC903-treated skin were excluded due to low quality of cDNA. Differential expression analysis was performed using the edgeR package (version 3.28.1) in R (version 3.6.2) and differentially expressed genes (DEGs) were defined as those with false discovery rate (*FDR*) < 0.05. The data discussed in this publication have been deposited in Gene Expression Omnibus database (accession number GSE180541).

For the comparison of gene expressions in skin MC populations (rMCs and integrinβ7^+^ MCs in NT and MC903-treated skin) with progenitor cells in BM (pro-GMPs and pre-GMPs), the published RNA-seq data (GSE132122) was used. The sequencing data were mapped in the same way as MC and logarithmically transformed RPKM (Reads Per Kilobase transcript, per Million mapped reads) of genes were used for Spearman Rank correlation after removing low expression genes (genes for which counts were greater than 10 for 70% of the samples were retained). Each of the study samples were correlated against all the samples in the published data and the correlation coefficient rho averaged for each of the cell type. This resulted in a single averaged correlation coefficient against each of the published cell type for each of the study sample. Kruskal-Wallis tests were used to test this averaged correlation coefficient rho between skin MC subsets for pro-GMPs and pre-GMPs to determine if there were significant differences.

### Intradermal-diphtheria toxin (DT) treatment to Mas-TRECK mice

Dorsal and ventral side ears of either Mas-TRECK mice or wild type (C57BL/6) mice (WT) were injected intradermally (i.d.) with 10 ng of diphtheria toxin (DT) (Sigma-Aldrich) in 20 µl of PBS for one time. Ears were treated with MC903 4 days after i.d. injection.

### Wsh/Wsh-BM chimera mice

Recipient wild type C57BL/6 (WT) mice were lethally irradiated with 9.5 Gy by an X ray irradiation device and received 5×10^6^ BM cells from either Wsh/Wsh mice or WT mice intravenously through the tail vein. The chimeric mice were subjected to the indicated experiments over 12 weeks after the reconstitution.

### Hydroxyproline Colorimetric Assay

Hydroxyproline of ears was measured using Hydroxyproline Assay Kit (Sigma-Aldrich). All procedures were performed according to the manufacturer’s protocol. Briefly, ears were homogenized in distilled water. 100 μl of Hydrochloric acid (12N) was added to 100 μl of homogenized solution, then samples were hydrolyzed by incubating at 120°C for 3 hours. Samples were vortexed and centrifuged at 10000g for 3 minutes, then 10 μl of the supernatant was transferred to a 96-well plate and evaporate in 60°C incubator. Add 100 μl of chloramine T reagent to each well and incubate for 5 minutes at room temperature. Then, add 100 μl of DMAB reagent and incubate for 90 minutes at 60°C. The absorbance at 560 nm was measured with a microplate reader (FlexStation 3).

### Adoptive transfer of BM cells to C57BL/6-*KitW_-sh_/W_-sh_* mice and C48/80 test

C57BL/6-*KitW_-sh_/W_-sh_* mice received 1×10^7^ BM cells from WT mice intravenously through the tail vein. The recipient mice were treated with MC903 over 8 weeks after the reconstitution. MC903 was applied to both ears for 10 days, then 1 μg of compound 48/80 (C48/80) (Sigma-Aldrich) in saline was intradermally injected on day 35.

### Immunostaining of human skin samples

Normal skin samples were obtained from non-involved parts of the triangular end of the surgically removed benign tumors. Lesional skin samples were obtained from skin biopsies of patients clinically diagnosed with AD. This study was approved by the ethics committee of the Kyoto University Graduate School of Medicine (R0743) and written informed consent was obtained from individuals and was conducted according to the principles of the Declaration of Helsinki. Paraffin-embedded skin samples were deparaffinized and subjected to heat-mediated antigen retrieval. Triple immunofluorescent staining was performed for c-kit, FcεRI, and integrinβ7. The antibodies used were anti-human CD117 (EP10; Leica Biosystems, Buffalo Grove, IL), anti-human FcεRI (9E1; Abcam), and biotin anti-mouse integrinβ7 (FIB504; Abcam) for the primary antibodies, and Alexa Fluor 647 anti-rabbit IgG antibody, Alexa Fluor 488 anti-mouse IgG antibody, and Alexa Fluor 594 streptavidin (Invitrogen) for the second antibodies. Coverslips were mounted with Prolong Gold Antifade Mountant with DAPI (ThermoFisher Scientific, Waltham, MA). Fluorescent images were analyzed with a Nikon confocal microscope (Nikon A1RMP, Nikon, Tokyo, Japan).

### Statistical analysis

Data are presented as the mean values ± standard deviation (SD). Statistical analyses were performed using GraphPad Prism (GraphPad Software, Inc., La Jolla, CA). Normal distribution was assumed for all samples. Unless indicated otherwise, a parametric Student’s *t*-test, a one-way ANOVA test, or a two-way ANOVA was used for comparing groups. A value of *P* < 0.05 at 95% confidence intervals was considered to indicate statistical significance and indicated with an asterisk. NS indicates not significant.

## Acknowledgments

We thank Dr. Masato Kubo for providing Mas-TRECK mice, and Ms Hiromi Doi and Ms Kaori Tomari for technical support. We also thank the Center for Anatomical, Pathological and Forensic Medical Research, Kyoto University Graduate School of Medicine, for preparing microscope slides. This work was supported by grants from the Japan Society for the Promotion of Science KAKENHI (JP15K09766, JP15H05096 [T.H.], 15H05790, 15H1155, 15K15417, 20H05697 [K.K.], 17J10049, and 21J40104), Japan Agency for Medical Research and Development (AMED) (16ek0410011h0003, 16he0902003h0002, JP20gm1210006 [K.K.]).

## Supplemental Information

Supplemental Information includes 5 figures.

## Supplemental Figure

**Figure S1.**
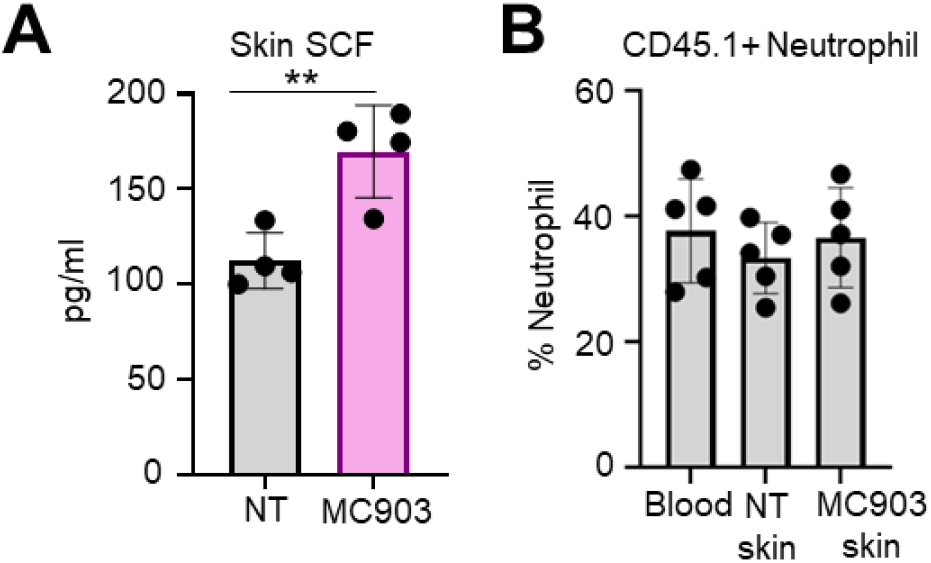
**A,** ELISA analysis of SCF levels in NT and MC903-treated skin (n = 4). **B,** The ratio of CD45.1^+^ neutrophils in blood, NT skin, and MC903-treated skin of WT CD45.2^+^ mice. (n = 5) Results are expressed as the mean ± standard deviation (SD). Data are representative of at least two independent experiments. **, p < 0.01

**Figure S2.**
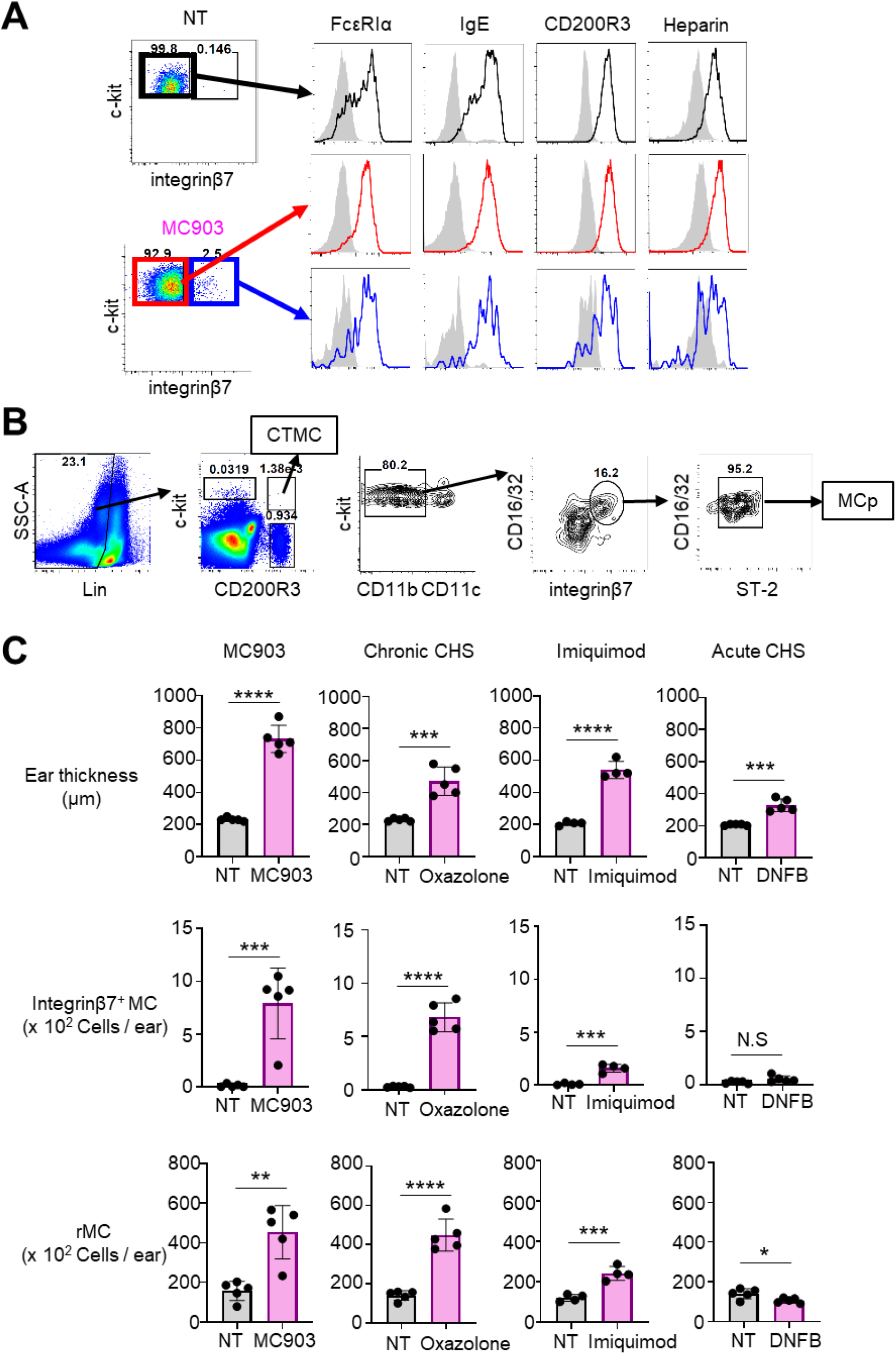
**A,** Representative flow cytometry panels for the expression of FcεRIα, and IgE, CD200R3, and heparin on integrinβ7^-^ MCs (rMCs) in NT skin (black line) and integrinβ7^-^ MCs (rMCs) (red line) and integrinβ7^+^ MCs (blue line) in MC903-trated skin. The gray area represents isotype control antibody. **B,** Gating strategy of CTMCs and MCps in blood. **C,** Comparison of ear thickness and the numbers of integrinβ7^+^ MCs and rMCs in the skin of NT and different dermatitis models including MC903-induced AD model (MC903) (n = 5), oxazolone-induced chronic contact hypersensitivity (chronic CHS) (n = 5), imiquimod-induced psoriasis-like dermatitis model (imiquimod) (n = 4), and DNFB-induced acute contact hypersensitivity (acute CHS) (n = 5). Results are expressed as the mean ±SD. Data are representative of at least two independent experiments. *, p < 0.05, **, p < 0.01, ***, p < 0.001, and ****, p < 0.0001. NS, not significant.

**Figure S3.**
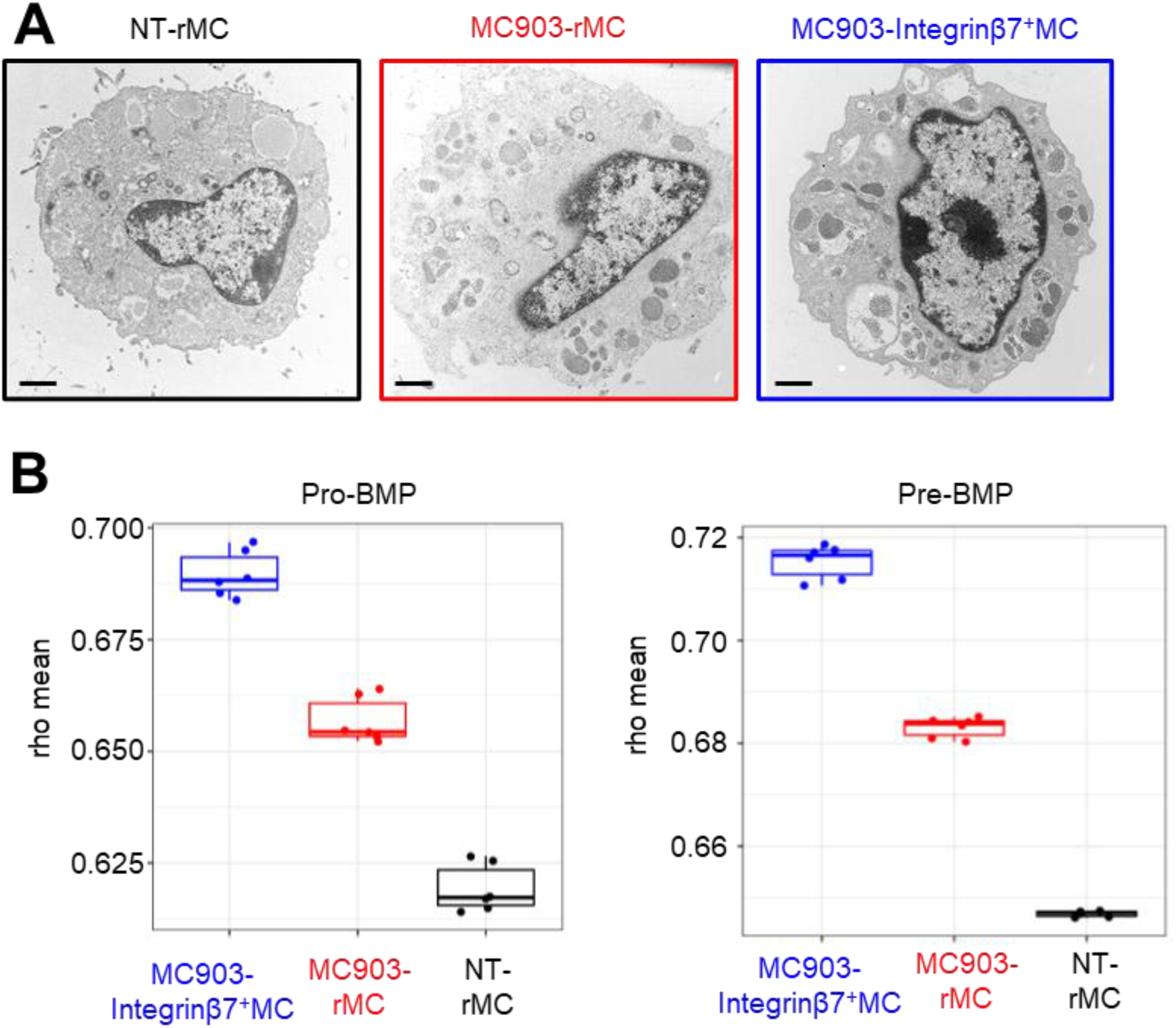
**A,** Electron micrographs of rMCs and integrinβ7^+^ MCs in NT and MC903-treated skin. Scale bar, 1 μm. **B,** Boxplots of averaged Spearman Rank correlation coefficient rho of the integrinβ7^+^ MCs in NT and MC903-treated skin sample gene expression against the Pro-BMPs (left) and Pre-BMPs (right). Kruskal-Wallis tests showed that there were significant differences between rMCs and integrinβ7^+^ MCs in NT and MC903-treated skin.

**Figure S4.**
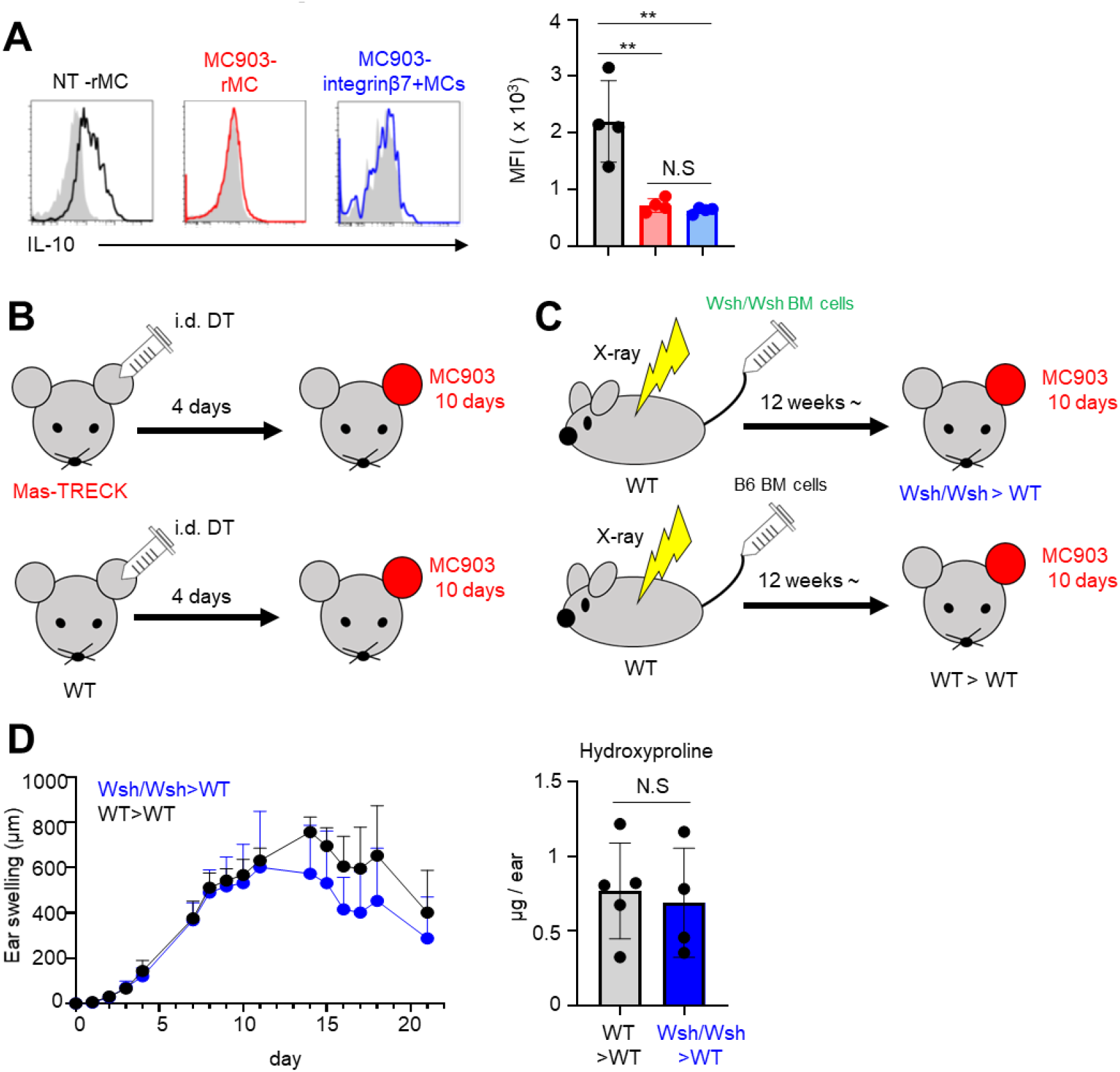
**A,** Representative flow cytometry panels for the expression of IL-10 on rMCs in NT skin (black line) and rMCs (red line) and integrinβ7^+^ MCs (blue line) in MC903-trated skin. The gray area represents isotype control antibody. Mean fluorescence intensity (MFI) of IL-10 was shown as a bar graph. (n = 4) **B,** The experimental scheme of MC903-induced AD model with/without resident MCs using Mas-TRECK mice. **C,** The experimental scheme of MC903-induced AD model with/without BM-derived MCs using Wsh/Wsh BM-chimera mice. **D,** Ear swelling of MC903-treated skin with/without BM-derived MCs until day 21 (left panel). Hydroxyproline level of WT>WT mice and Wsh/Wsh>WT mice (right panel). (n = 4 or 5) Results are expressed as the mean ±SD. Data are representative of at least two independent experiments. **, p < 0.01, NS, not significant.

**Figure S5.**
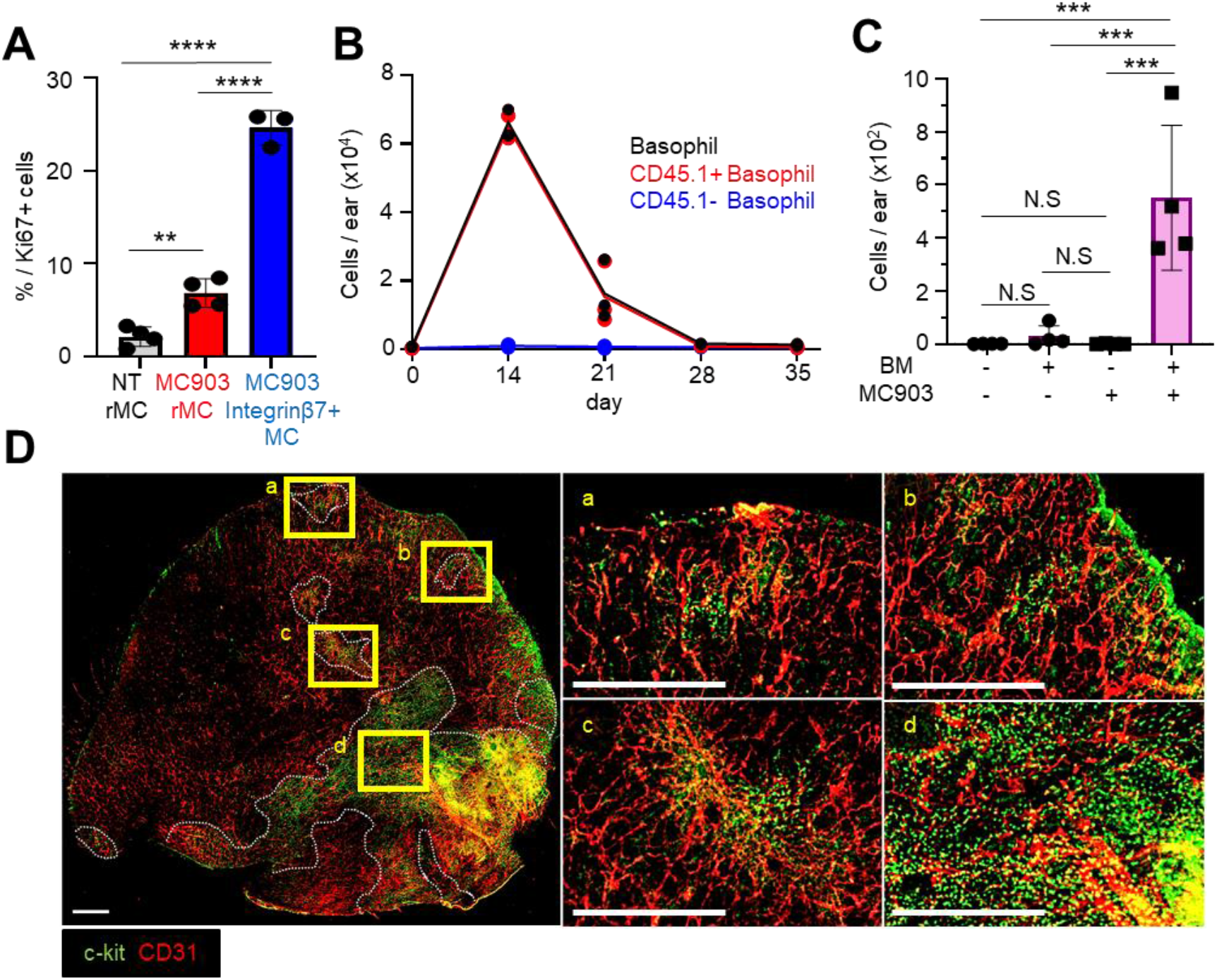
**A,** The percentage of Ki67-positive cells in rMCs and integrinβ7^+^ MCs in NT and/or MC903-treated skin. (n = 3 or 4) **B,** The numbers of basophils (Live^+^ CD45^+^ CD11b^-^ CD200R3^+^ c-kit^-^ cells), CD45.1^+^ basophils, and CD45,1^-^ basophils in the skin on day 0, 14, 21, 28, and 35 in MC903-induced AD model. (n = 2 - 3) **C,** The numbers of skin MCs in NT or MC903-treated skin of Wsh/Wsh mice ears with or without adoptive transfer of BM cells. (n = 4) **C,** Whole mount immunostaining of c-kit (green) and CD31 (red) in ear skin. Scale bar, 1 mm (left panel). Dotted white line showed the gathering of c-kit positive cells. Results are expressed as the mean ±SD. Data are representative of at least two independent experiments. ***, p < 0.001. NS, not significant.

